# CDK-regulated phase separation seeded by histone genes ensures precise growth and function of Histone Locus Bodies

**DOI:** 10.1101/789933

**Authors:** Woonyung Hur, Marco Tarzia, Victoria E. Deneke, Esteban A. Terzo, Robert J. Duronio, Stefano Di Talia

## Abstract

Many membrane-less organelles form through liquid-liquid phase separation, but how their size is controlled and whether size is linked to function remain poorly understood. The Histone Locus Body (HLB) is an evolutionarily conserved nuclear body that regulates the transcription and processing of histone mRNAs. Here, we show that *Drosophila* HLBs form through phase separation of the scaffold protein multi-sex combs (Mxc). The size of HLBs is controlled in a precise and dynamic manner that is dependent on the cell cycle and zygotic gene activation. Control of HLB growth is achieved by a mechanism integrating nascent mRNAs at the histone locus, which catalyzes phase separation, and the nuclear concentration of Mxc, which is controlled by the activity of cyclin-dependent kinases. Reduced Cdk2 activity results in smaller HLBs and the appearance of nascent, misprocessed histone mRNAs. Our experiments thus identify a mechanism linking nuclear body growth and size with gene expression.

## INTRODUCTION

Many nuclear bodies form by liquid-liquid phase separation (Berry et al., 2018; Brangwynne, 2013; Brangwynne et al., 2011; Falahati et al., 2016; Falahati and Wieschaus, 2017; Hyman et al., 2014; Li et al., 2012; Mitrea and Kriwacki, 2016), the physical process by which certain fluids spontaneously separate into stable regions of different concentration or chemical composition. A major function of phase separation is to concentrate components to enhance the rate of biochemical reactions (Stroberg and Schnell, 2018). Consequently, the size of nuclear bodies should determine their overall biochemical activity. Yet, it remains unclear whether there are molecular and physical mechanisms that ensure precise regulation of the growth and size of nuclear bodies. Since phase separation is an intrinsically stochastic process, understanding whether the size of nuclear bodies or other types of biomolecular condensates is linked to their function and which mechanisms might ensure accurate size control remain fundamental open questions. Here, we study the molecular and physical mechanisms of formation of the Histone Locus Body (HLB), an evolutionarily conserved nuclear body that plays a fundamental role in the transcription and processing of histone mRNAs (Duronio and Marzluff, 2017). We focus on the syncytial nuclear cycles that lead to the maternal-to-zygotic transition in the *Drosophila* embryo, a stage of development when we can precisely examine whether HLBs form by phase separation and test how phase separation might be affected by the cell cycle and activation of zygotic gene expression.

The HLB forms at replication-dependent (RD) histone genes; i.e. histone genes expressed only during the S phase of the cell cycle (Duronio and Marzluff, 2017; Marzluff and Duronio, 2002). In *Drosophila*, these genes are located at a single locus containing a tandem array of about one hundred gene copies (Figure 1A) (Bongartz and Schloissnig, 2019; Lifton et al., 1978; McKay et al., 2015). The nascent histone transcripts are processed via a single endonucleolytic cleavage, resulting in mRNAs with a unique 3’ end that is not polyadenylated (Marzluff and Koreski, 2017). The HLB contains components necessary for both the transcription and specialized processing of these mRNAs (Burch et al., 2011; Duronio and Marzluff, 2017; Mao et al., 2011; Matera et al., 2009; Tatomer et al., 2016; White et al., 2011). Reciprocally, transcription of RD histone genes contributes to HLB formation (Salzler et al., 2013; Shevtsov and Dundr, 2011). Failure to localize FLASH, an essential histone pre-mRNA processing factor, to the HLB reduces the efficiency of histone pre-mRNA processing, suggesting that concentrating proteins within the HLB is necessary for histone mRNA biogenesis (Tatomer et al., 2016). Thus, understanding the processes regulating HLB assembly and organization will provide insight into how biomolecular condensates contribute to critical cellular functions like gene expression (Sawyer et al., 2019).

**Figure 1.**
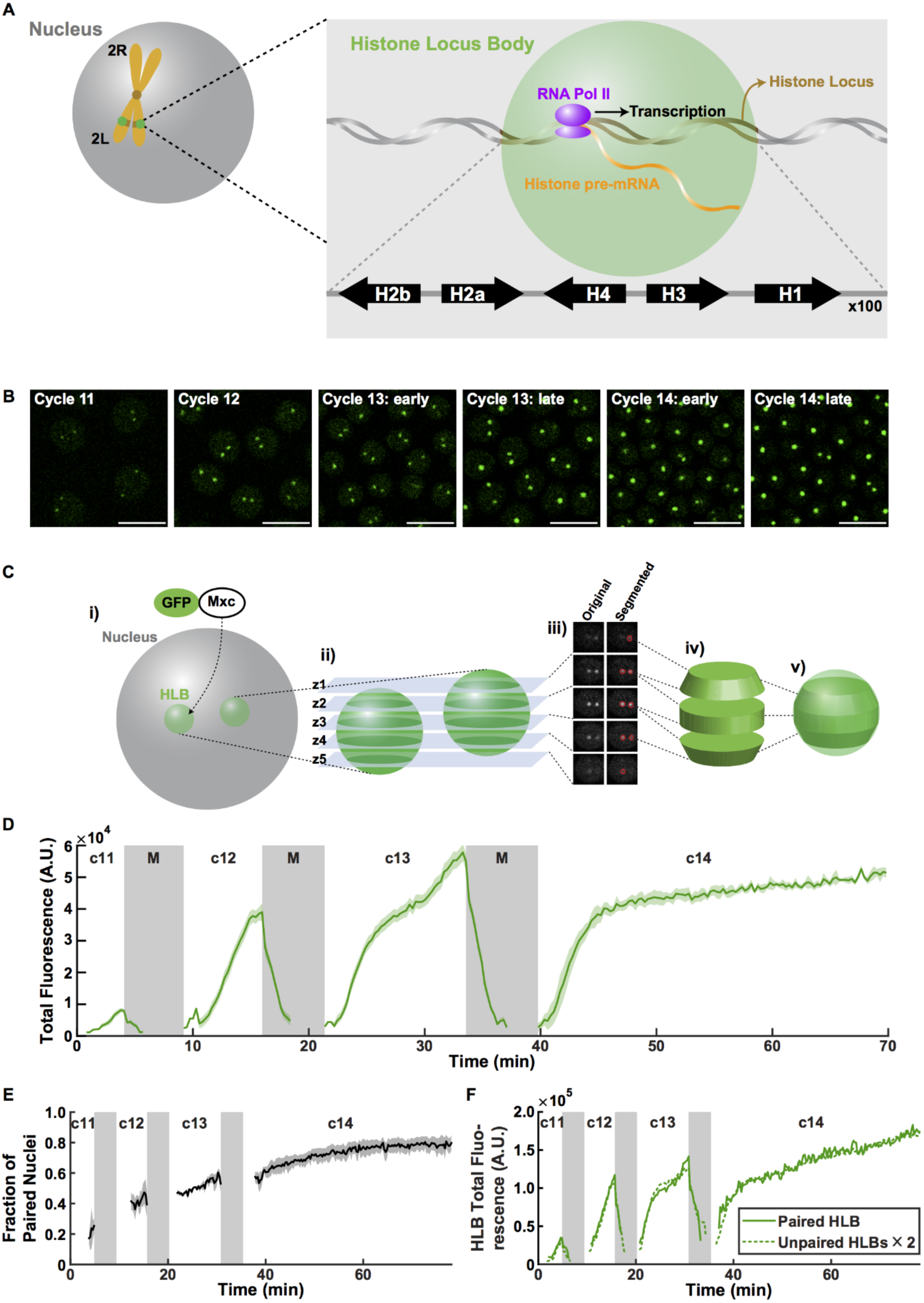
Cell cycle-dependent regulation of HLB size. (A) Schematic of the Histone Locus Body and the *Drosophila* histone locus located on the left arm of the second chromosome (2L). (B) Representative images of HLBs (GFP-Mxc) during S phase of syncytial cycles 11-14. HLBs show homogeneous size distribution dependent on cycle stage. Scale bars, 10µm. c, cycle. (C) Schematic of HLB size quantification. i) HLBs were visualized by expressing a GFP-Mxc fusion protein in the presence of endogenous Mxc. ii) Multiple z stack images were acquired with confocal microscopy. iii) HLBs were segmented using threshold-based computational segmentation. iv) HLB size was approximated by measuring the trapezoidal cone shaped space occupied by multiple HLB masks. v) HLB ‘Total Fluorescence’ was quantified by multiplying the approximated size by the average fluorescence intensity level of the HLB masks. (D) HLB total fluorescence as a function of time for cycles 11-14. Shaded green area, SEM. (E) Fraction of nuclei with paired chromosomes over time. (F) Comparison between the average Mxc total fluorescence of fused HLBs (‘paired (fused) HLB’) and that of two HLBs that have not fused (‘unpaired HLBs multiplied by 2’). c, cycle. M, mitosis. A.U., arbitrary units.

A number of lines of inquiry have indicated that both hierarchical and stochastic recruitment of factors contribute to HLB formation (Shevtsov and Dundr, 2011; White et al., 2011). For instance, confocal microscopy of fixed syncytial fly embryos revealed that some HLB components arrive at the histone locus prior to others (White et al., 2011). In these embryos HLBs begin assembling at the onset of zygotic RD histone transcription, but the relationship between HLB assembly and transcription is unclear. Moreover, if and when the HLB demonstrates properties of phase separation have not been experimentally determined, and whether phase separation contributes to histone mRNA biosynthesis is unknown.

A key mediator of HLB assembly in *Drosophila* is the product of the *multi sex combs* gene (*mxc*). Mxc is the *Drosophila* orthologue of human NPAT, one of the first identified cyclin E/Cdk2 substrates (Zhao et al., 1998). Both NPAT and Mxc accumulate only in the HLB and provide the most definitive marker for HLB formation within cells (Ma et al., 2000; White et al., 2011; Zhao et al., 2000). Mxc/NPAT is likely the main scaffolding protein for the HLB, and in its absence the HLB does not form and RD histone genes are not expressed (Wei et al., 2003; White et al., 2011; Ye et al., 2003). Like NPAT, Mxc is a large (1837 amino acids) protein composed mostly of predicted intrinsically disordered regions, with an N-terminal LisH domain that facilitates oligomerization and is essential for Mxc function *in vivo* (Terzo et al., 2015; Wei et al., 2003). Because Mxc also is a Cyclin E/Cdk2 target (White et al., 2011), it and NPAT likely function to integrate HLB formation with cell cycle progression, although how this occurs is unknown.

To gain insight into the mechanisms of HLBs assembly and size control, we used confocal microscopy to image the dynamics of HLB formation in living *Drosophila* embryos expressing a GFP-Mxc fusion protein (Figure 1B) (Terzo et al., 2015; White et al., 2011). By combining this live imaging approach with genetic manipulation and quantitative analyses, we show that HLBs form through phase separation. In addition, we discovered a role for cyclin dependent kinase activity in HLB size control, and show that regulation of HLB size is important for normal processing of RD histone mRNAs.

## RESULTS

### HLBs undergo precise growth dynamics linked to histone biogenesis

In order to quantify HLB dynamics, we developed a computational method to automatically segment nuclei and GFP-Mxc-labeled HLBs within time lapsed confocal images (Figures 1C and 1D; Methods). In early *Drosophila* embryos, HLBs form during S phase and dissolve during mitosis, confirming cell cycle dependent dynamics (Video S1). Since HLBs form at the histone locus (Salzler et al., 2013), two HLBs often can be observed on the two homologous chromosomes in early embryos. The two HLBs come together upon homologous chromosome pairing, and the proportion of nuclei with paired homologs increases throughout the syncytial blastoderm stage (Hiraoka et al., 1993) (Figure 1E). We measured the total GFP-Mxc fluorescence of paired and unpaired HLBs and confirmed that paired HLBs contained twice as much Mxc molecules as unpaired HLBs (Figure 1F), indicating that chromosome pairing simply merged two HLBs together but did not promote further growth. Since the total GFP-Mxc fluorescence of HLBs in each nucleus is unaffected by chromosome pairing, we used this quantity to describe HLB size during the syncytial blastoderm stage (cycles 11-14). These analyses revealed highly reproducible HLB dynamics (Figure 1D). Similar results were obtained using the volume of HLBs in each nucleus (Figure S1A). As previously reported (White et al., 2007), HLBs are first detected during cycle 11 and disappear during mitosis before growing to a significant size. During cycles 12 and 13, S phase slows down and HLBs become significantly larger. The larger HLB size observed during cycles 12 and 13 was not simply due to a longer S phase, as the HLBs grow at a faster rate in these cycles (Figures S1B-S1C). The more rapid growth of HLBs during cycles 12-13 than cycle 11 is also observed in embryos mutant for the DNA replication checkpoint kinases Chk1 and Chk2 (*grapes, loki*), which unlike wild type embryos have roughly the same S phase durations during cycles 11-13 (Figures S1B-S1C). Thus, our results indicated that inputs other than duration of S phase contribute to the rate of growth of HLBs.

In principle, the formation of nuclear bodies by liquid-liquid phase separation could be influenced by an initial stochastic nucleation step, which would result in highly variable HLB size. To determine whether the mechanisms of HLB formation are stochastic or able to ensure consistent growth dynamics across the embryo, we measured HLB size distribution in individual nuclei. We found that the distribution of total GFP-Mxc fluorescence within HLBs is approximately Gaussian with a low coefficient of variation (0.2-0.3) (Figure 2A-2C). Moreover, the variability of HLB sizes is low throughout the syncytial blastoderm stage (Figure 2C), indicating that HLB growth is highly reproducible and uniform across all the nuclei. We hypothesized that such precise HLB growth and size control might play a role in the regulation of histone biogenesis during development. To test the possible functional relationship between HLB size and histone mRNA biosynthesis, we measured levels of nascent H3 histone mRNA by immuno-FISH (Figure 2D; Methods) and HLB size using the monoclonal antibody MPM-2, which specifically binds phosphorylated Mxc (Figure 2E) (White et al., 2011; White et al., 2007). Immunofluorescence experiments showed that the levels of total and phosphorylated Mxc closely correlate in interphase (Figure S2A), indicating that these two different parameters can be used interchangeably to measure HLB size. We found a significant correlation between the amount of nascent H3 mRNA and phosphorylated Mxc present in the HLB (Figure 2F), suggesting that nascent histone mRNA and HLB size are co-regulated. Thus, we set out to dissect the molecular mechanisms that ensure precise control of HLB growth dynamics during early embryogenesis.

**Figure 2.**
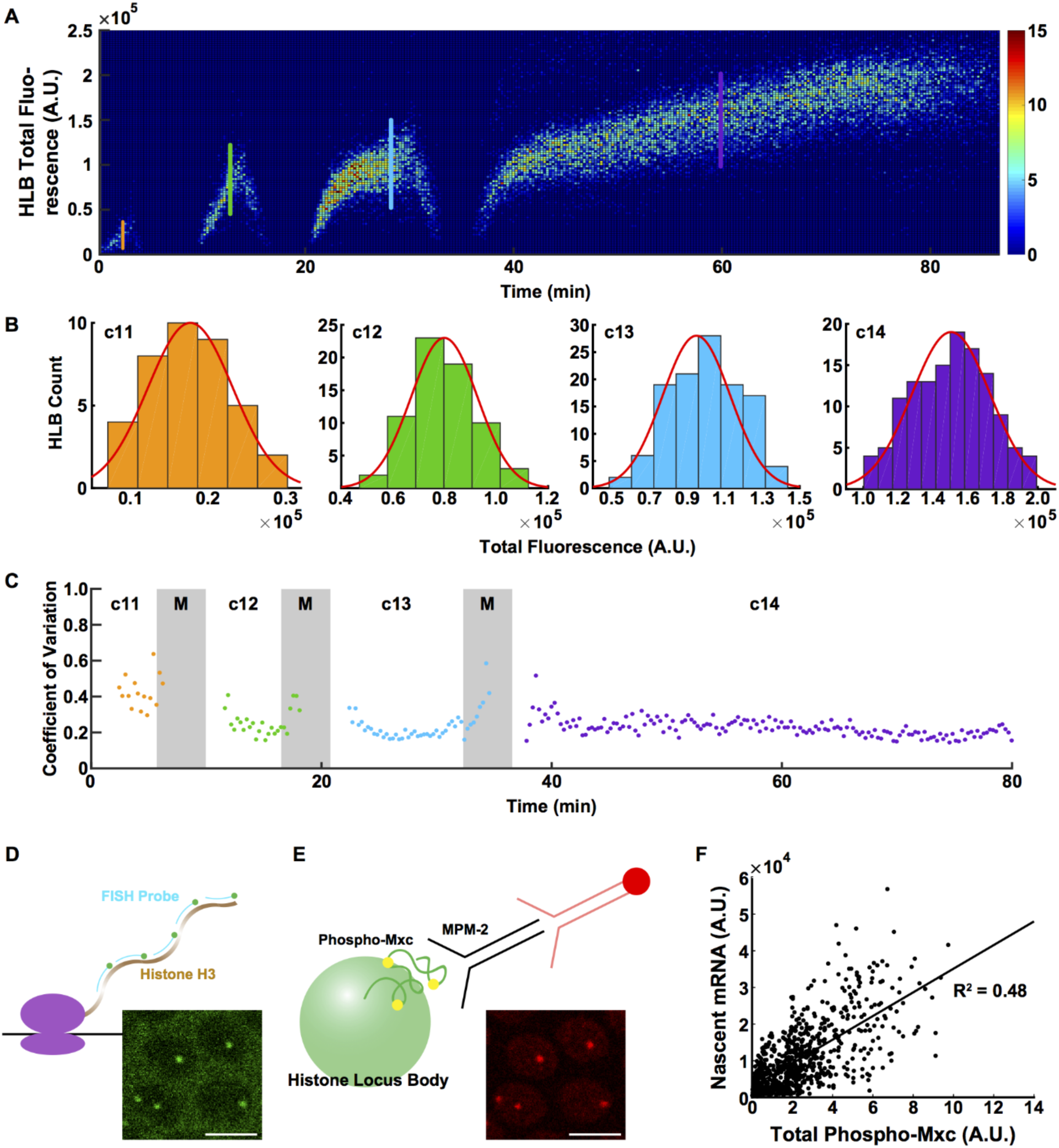
Precise HLB growth control is linked to histone mRNA biogenesis. (A) 3D histogram of HLB total fluorescence distribution. (B) HLB total fluorescence distribution for each cell cycle during the syncytial blastoderm stage. Each histogram indicates the total fluorescence distribution at the time points marked with the line with the same color in Figure 2A. Red line, fitted normal distribution. (C) Coefficient of variation (CoV) for HLB total fluorescence over the syncytial blastoderm stage. (D) Diagram of FISH (fluorescence in situ hybridization) probes of nascent histone mRNAs and a representative image of an embryo stained with probes targeting the coding sequence of histone H3. (E) Diagram of monoclonal antibody MPM-2, which recognizes phosphorylated Mxc within the HLB. Staining with histone H3 probe and MPM-2 antibody showed high colocalization. (F) Correlation of histone H3 nascent mRNA levels and total phosphorylated Mxc levels. A.U., arbitrary units. c, cycle. M, mitosis. Scale bars, 5µm.

### HLBs form by phase separation

We reasoned that, similar to other NBs, the behavior of the HLB might be governed by liquid-liquid phase separation (Brangwynne, 2013; Dundr, 2012; Hyman et al., 2014; Mao et al., 2011; Matera et al., 2009; Mitrea and Kriwacki, 2016). Taking advantage of the fact that homologous chromosomes actively pair during nuclear divisions in early *Drosophila* embryos (Hiraoka et al., 1993), we imaged individual HLBs as they become closely juxtaposed and then fuse (Figure 3A; Video S2). Since phase separated droplets tend to be spherical, we tested whether they recover a spherical shape following fusion. We quantified the circularity index of HLBs before, during, and after fusion to address if they behave as liquid droplets (Methods). The circularity index dropped significantly as HLBs came in contact with each other but quickly recovered within less than a minute from initial contact as the two HLBs fused into one (Figure 3B). This observation indicates that HLBs behave similarly to liquid droplets (Freeman Rosenzweig et al., 2017; Strom et al., 2017). The liquid nature of the HLB was further supported by Fluorescence Recovery After Photobleaching (FRAP) experiments (Figure 3C), demonstrating that after bleaching GFP-Mxc-labelled HLBs recover fluorescence on timescales of seconds to minutes (Figure 3D and 3F). The time of recovery could be estimated from a linear fit of the logarithm of the difference of fluorescence intensities between the unbleached and the bleached HLBs (Figure 3E). This analysis demonstrates that the dwell-time of Mxc in HLBs increases rapidly as a function of the size of the droplet (Figure 3F), which is consistent with a phase separation scenario (Huang et al., 2019). Phase separation is also supported by the observation that Mxc fails to accumulate at the histone locus when a mutation is introduced in the LisH domain (Mxc^LisH-AAA^), which mediates protein-protein self-interaction and Mxc oligomerization (Terzo et al., 2015) (Figure S4A).

**Figure 3.**
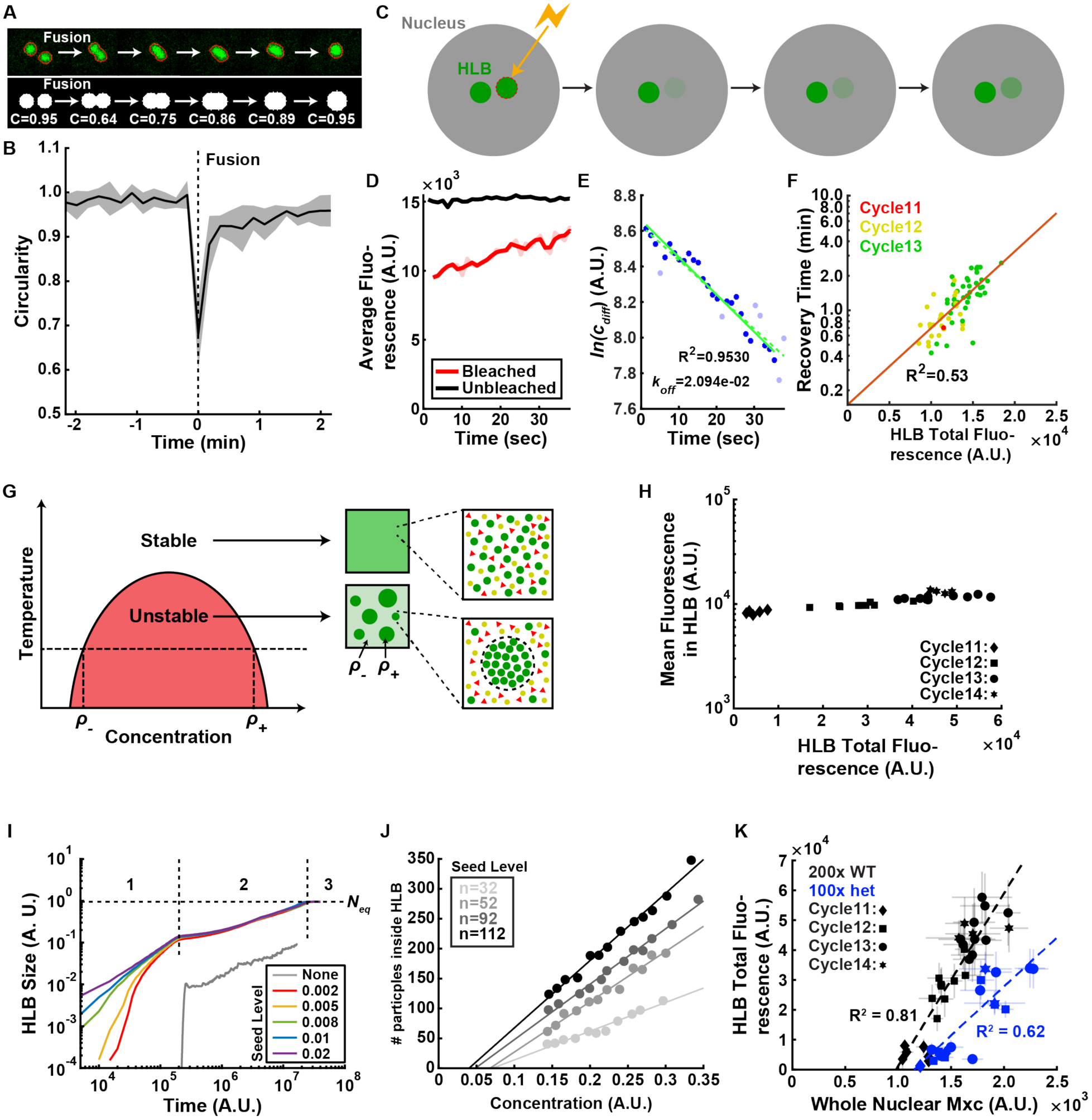
The HLB forms by phase separation. (A) Top: Example of HLB fusion upon chromosome pairing during S phase in *Drosophila* embryos. Bottom: Simulation of HLB fusion event and quantification of droplet circularity during the fusion. (B) Quantification of HLB circularity index during fusion events. Multiple quantifications were performed, aligned by the fusion timing (t=0), and averaged. Shaded area, SEM. (C) Schematic of HLB FRAP experiment. (D) Example of unbleached (black) versus bleached (red) HLB fluorescence level quantification. Dim red line indicates original data and the solid red line indicates the same data after noise correction, which is used for analysis in panel (E). (E) Logarithm of the difference between bleached (red in (D)) and unbleached (black in (D)) HLB fluorescence from (D) over time. The initial fit (dashed green line) was generated from all data points (blue and light blue points). From this initial fit, points whose residuals were greater than 2 standard deviations were removed from the dataset (light blue points). The best fit line was then computed on the dataset lacking the outliers (solid green line). The export rate *k*_*off*_ was measured from the slope of the solid green line. (F) HLB recovery time (on logscale) as a function of the HLB total fluorescence. (G) Phase diagram for a liquid-liquid phase separation. At a given temperature and concentration, a mixture of macromolecules with the potential to undergo liquid-liquid phase separation becomes thermodynamically unstable and separates into two different phases with concentrations of *ρ*_*-*_ and *ρ*_*+*_. Red area indicates the portion of the phase diagram in which phase separation occurs. Thus, *ρ*_*-*_ and *ρ*_*+*_ can be determined for any given temperature (horizontal dotted line). Red triangles and yellow circles represent other soluble proteins that do not participate in the phase separation; while green circles are the phase-separated component (e.g. GFP-Mxc). Adapted from (Brangwynne, 2013). (H) Quantification of average Mxc concentration inside HLB versus HLB total fluorescence. (I) Numerical solution of Cahn-Hilliard (CH) equation in 2D lattice reveals that HLB growth proceeds in three stages: 1) seeding level-dependent fast growth, 2) slower growth independent of the seeding level, and 3) equilibrium phase where HLB size is only dependent on nuclear Mxc concentration. (J) A particle-based simulation of HLB growth in the 3D lattice shows that HLB size (y-axis) depends on the seeding level (shades of gray) and Mxc concentration (x-axis) in the first regime indicated as “1” in (I). (K) Experimental data of HLB total fluorescence inside each nucleus and nuclear Mxc concentration, for WT (∼200 histone repeats) and histone deficiency heterozygote mutant (∼100 histone repeats). A.U., arbitrary units.

### Theoretical analysis reveals a mechanism for the precise control of HLB growth and size

The theory of phase separation indicates that within a particular range of temperatures and concentrations certain mixed liquids are thermodynamically unstable (Brangwynne, 2013). Consequently, these liquids spontaneously separate in two phases of different concentration, indicated respectively as ρ- and ρ+ for a given temperature (Figure 3G). We can model the nucleoplasm as a mixed liquid that separates into a phase of high Mxc concentration ρ_+_ (the HLB), and a phase of low Mxc concentration ρ_-_. Theory predicts that the concentrations ρ_+_ and ρ_-_ do not depend on the overall nuclear concentration of Mxc and HLB size. This prediction was confirmed experimentally for ρ_+_ (Figure 3H), arguing that phase separation ensures that the Mxc concentration within the HLB is reproducibly controlled. For a phase separation process at thermodynamic equilibrium, the size of the HLB should depend solely on the nuclear concentration of Mxc, a theoretical prediction not supported by published observations that the histone locus is required for proper formation of the HLB (Salzler et al., 2013). Thus, to further dissect how HLBs form, we used numerical simulations of a standard theoretical model (Cahn-Hilliard equation) describing phase separation (Bray, 2002) and previously used to describe the formation of nuclear bodies (Berry et al., 2015). We introduced a modification in the model to include the role of the histone locus and nascent histone mRNA in facilitating the formation of HLBs (Methods). To this end, we imposed that only the state of high concentration is stable in the immediate vicinity of the histone locus. Hereafter, we refer to this as seeding. Different levels of seeding were implemented by changing the volume of the region where seeding occurs. Simulations indicate that phase separation initiated by seeding can ensure precise control of HLB growth (Figure S3A) and proceeds in three stages (Figure 3I; Methods). In the first stage, the growth of the HLB is rapid and strongly influenced by the level of seeding. In the second phase, growth slows down and becomes largely independent of the seeding level. In the third phase, the system reaches thermodynamic equilibrium with the HLB size acquiring an asymptotic value. This equilibrium size is dictated only by the nuclear Mxc concentration and is independent of seeding. A more realistic particle-based model of phase separation (Methods) confirmed that, in the first regime, the size of the HLB is influenced in a quantitative manner by the level of seeding (Figure 3J). We derived a relationship between HLB size and the concentration of Mxc, delineated as a straight line passing through an x-intercept with the slope controlled by the level of seeding (Figure 3J). Our observational data from embryos confirmed these theoretical predictions. For instance, we demonstrated the dependency of the slope on seeding levels by comparing data from wild type embryos to data from embryos heterozygous for a deficiency covering the entire histone locus (i.e. containing half the number of seeding histone genes, Figure 3K). Collectively, these results suggest that there is a non-equilibrium regime in the formation of phase-separated droplets, where both the levels of seeding and concentration play a role in determining droplet size. This regime differs from the behavior at equilibrium when the size of the droplets depends solely on concentration and is independent of seeding.

### Nascent histone mRNAs provide a seed for HLB formation and influence HLB growth

To strengthen the previous conclusion, we set out to further test the role of seeding in the regulation of HLB growth by quantifying the dynamics of GFP-Mxc foci formation in embryos homozygous for a deletion of the entire histone locus. The maternal supply of histones allows these embryos to develop normally until gastrulation but they lack any seeding activity because of the absence of histone genes. Our simulations predicted that in this scenario a variable number of droplets of much smaller and heterogeneous size should form (Figure 3I; Methods). We found that GFP-Mxc foci are significantly smaller when the histone locus is absent (Figure 4A and 4B; Video S3), a behavior that is expected for phase-separated objects whose formation is initiated by a stochastic fluctuation (nucleation) rather than in a regulated manner through seeding. In the absence of seeding, small droplets are spontaneously formed by thermal fluctuations. Yet, they are associated with a high energetic cost due to surface tension and are almost always reabsorbed. Phase separation thus only initiates when a rare droplet, large enough to overcome the energetic barrier, is created. Conversely, in the presence of seeding, the association of Mxc with the histone locus lowers this energetic barrier, thereby allowing the formation of HLBs in a much shorter time compared to the time scale over which droplets start to form under spontaneous phase separation. Consistent with this physical interpretation, GFP-Mxc foci formed in embryos lacking the histone locus are variable in number (often more than 2 per nucleus form; Figure 4A), non-uniformly distributed in size (Figures S4B-S4E), and highly transient. During nuclear cycle 14, when S phase duration is significantly longer, some GFP-Mxc foci grow larger than in the previous cycles, as expected (Figure S4F). Taken together, our data suggest that the small bodies previously named proto-HLBs (Salzler et al., 2013) result from the intrinsic ability of Mxc to phase separate. Therefore, we conclude that HLBs form through a phase separation process and that in wild type embryos the intrinsic stochasticity associated with nucleation is supplanted by seeding from the histone locus to ensure dynamic and reproducible control of size and histone biogenesis.

**Figure 4.**
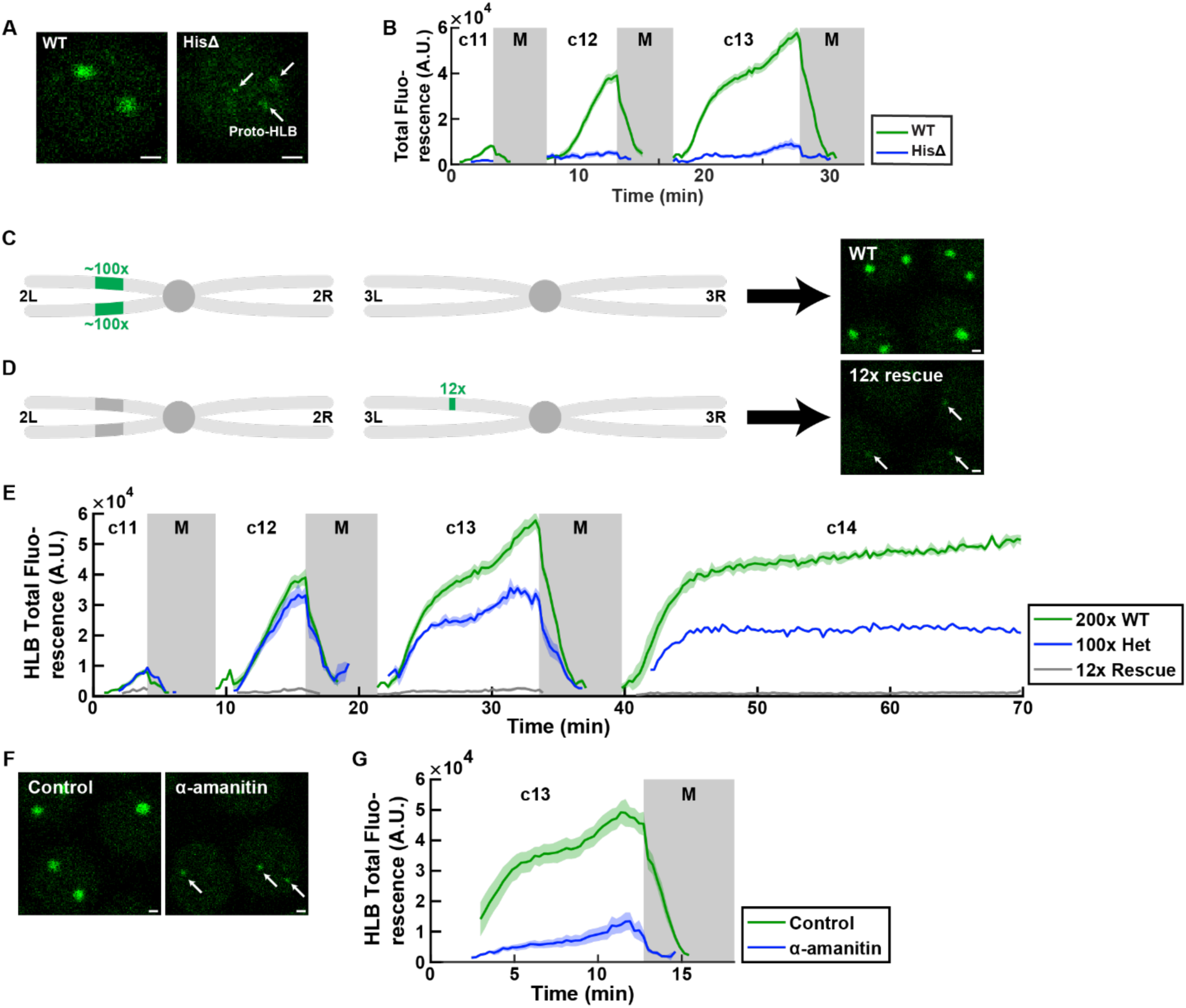
Proper HLB formation requires seeding from the histone locus. (A) Examples of WT HLB and the transient proto-HLBs observed in homozygous histone deficiency (HisΔ) embryos in cycle 13. Arrows indicate proto-HLBs. (B) Quantification of total fluorescence for WT and HisΔ embryos. (C) Schematic of the second and third chromosome for WT flies. Endogenous histone locus is on 2L, and it consists of about 100 copies of tandem repeats of all the canonical histone genes (H3, H4, H2a, H2b, and H1). (D) Schematic of the second and third chromosome for 12x histone rescue transgenic flies. 12 copies of histone repeats were inserted into chromosome 3L. Images on the right in (C) and (D) are confocal images of HLBs from each respective genotype in cycle 13. Arrows indicate HLBs. Images were acquired under identical imaging conditions. (E) HLB total fluorescence for various numbers of histone repeats (200x, 100x, and 12x). (F) HLBs in control and α-amanitin injected embryos. Arrows indicate HLBs. (G) HLB total fluorescence for DMSO control and α-amanitin injected embryos. Shaded area, SEM. A.U., arbitrary units. c, cycle. M, mitosis. Scale bars, 1µm.

To further test the role of the histone locus as a seed facilitating growth of the HLB, we addressed quantitatively how changes in the level of seeding affect HLB size. First, we compared the growth of HLBs in wild type embryos and embryos heterozygous for a deficiency deleting the entire histone locus. Our experiments show that, with the possible exception of cycle 12, the size of the HLB when only one histone locus is present is about half of the sum of the two HLB sizes in wild type (Figure 4E and Figure S4G). This observation indicates that HLB growth is influenced quantitatively by seeding at the histone locus and thus favors a model in which HLB formation is in the first regime, far from thermodynamic equilibrium (Figure 3I; Figure S3C; Methods). At equilibrium, the size of the HLB in embryos carrying only one copy of the histone locus would have been comparable to the sum of sizes of the two HLBs in wild type (Figure S3C; Methods). We also used an ectopic transgene containing 12 copies of histone repeats (Figure 4D) that can rescue deletion of the entire endogenous histone locus, which has about 100 histone gene repeats on each homologous chromosome (Figure 4C) (McKay et al., 2015). We observed that, as previously reported (McKay et al., 2015), embryos with 12 copies of histone repeats form much smaller HLBs than wild type (Figure 4E), further supporting a role for the level of seeding in the regulation of HLB growth.

Finally, we tested whether nascent mRNAs contribute to seeding. To this end, we injected embryos with alpha-amanitin, an inhibitor of Pol II transcription. We observed that alpha-amanitin injected embryos formed significantly smaller HLBs (Figures 4F and 4G), while retaining normal dynamics of the nuclear cycles and Mxc concentration in the nucleoplasm (Figures S4H-S4J). This result indicates that nascent mRNAs are an important catalyzer of HLB growth (Heyn et al., 2017). Since still only one or two HLBs are observed after alpha-amanitin treatment (presumably at the histone locus), and since HLBs can form at the histone locus in the G1 phase of the cell cycle when RD histone genes are not transcribed (White et al., 2011), we conclude that DNA at the histone locus can seed phase separation and that such seeding is strongly enhanced by nascent histone mRNA.

### Cdk activity controls HLB growth

To gain further insight into the molecular mechanisms of HLB formation, we investigated HLB cell cycle regulation. Previous reports have shown that NPAT/Mxc is a substrate of Cyclin E/Cdk2 and that its phosphorylated form was detected in HLBs (White et al., 2007; Zhao et al., 1998). In addition, Cyclin E/Cdk2 has been shown to stimulate NPAT/Mxc mediated activation of histone gene expression (Ma et al., 2000; White et al., 2011; White et al., 2007; Ye et al., 2003; Zhao et al., 1998; Zhao et al., 2000). Therefore, we hypothesized that Cyclin E/Cdk2 plays an important role in HLB size control. To test this hypothesis, we utilized SNS-032, a small molecule CDK inhibitor with higher specificity towards Cdk2 than other Cdks (Ali et al., 2009; Chen et al., 2009). We injected SNS-032 into embryos entering cycle 13 and visualized HLB dynamics. Since some Cdks are known to regulate transcription, we verified that drug injection did not affect zygotic expression of a gene not associated with the HLB (*sloppy paired*; Figures S5B-S5C). To infer the efficiency of the drug treatment, we performed these experiments in embryos expressing GFP-Mxc as well as a biosensor that is phosphorylated by several Cyclin/Cdk complexes (Schwarz et al., 2018; Spencer et al., 2013), including Cyclin E/Cdk2 (Figure 5A). The biosensor protein is excluded from the nucleus after phosphorylation by Cdks (Figure 5B and Video S4), and thus the ratio of cytoplasmic-to-nuclear sensor protein provides a readout of Cdk-mediated phosphorylation (Figure 5C). During the syncytial blastoderm cycles, the localization of the biosensor oscillates with nuclear exclusion increasing over each S phase and rapidly decreasing prior to mitotic entry (Figure 5D). Mutating the putative Cdk phosphorylation residues on the biosensor to alanine led to loss of nuclear exclusion (Figure S5D). The phosphorylation of the biosensor and that of Mxc (detected by the MPM-2 antibody) have very similar dynamics, suggesting that the biosensor is an accurate proxy for the phosphorylation status of S-phase Cdk targets (Figure S5F). Treatment with the SNS-032 Cdk inhibitor significantly reduced the activity readout from the sensor and suppressed HLB growth (Figure 5E; Video S5). While at low doses the drug did not perturb nuclear cycle timing, at high doses, the nuclear cycle lengthened, likely through inhibition of Cdk1 (Figure S5A). To quantify the relationship between HLB size and the phosphorylation of the sensor, we measured HLB size about 10 minutes into S phase after SNS-032 injection as well as the biosensor activation rate (Figure 5E). We found that lower activation rate resulted in smaller HLB size (Figure 5F), consistent with a Cdk contribution to HLB growth.

**Figure 5.**
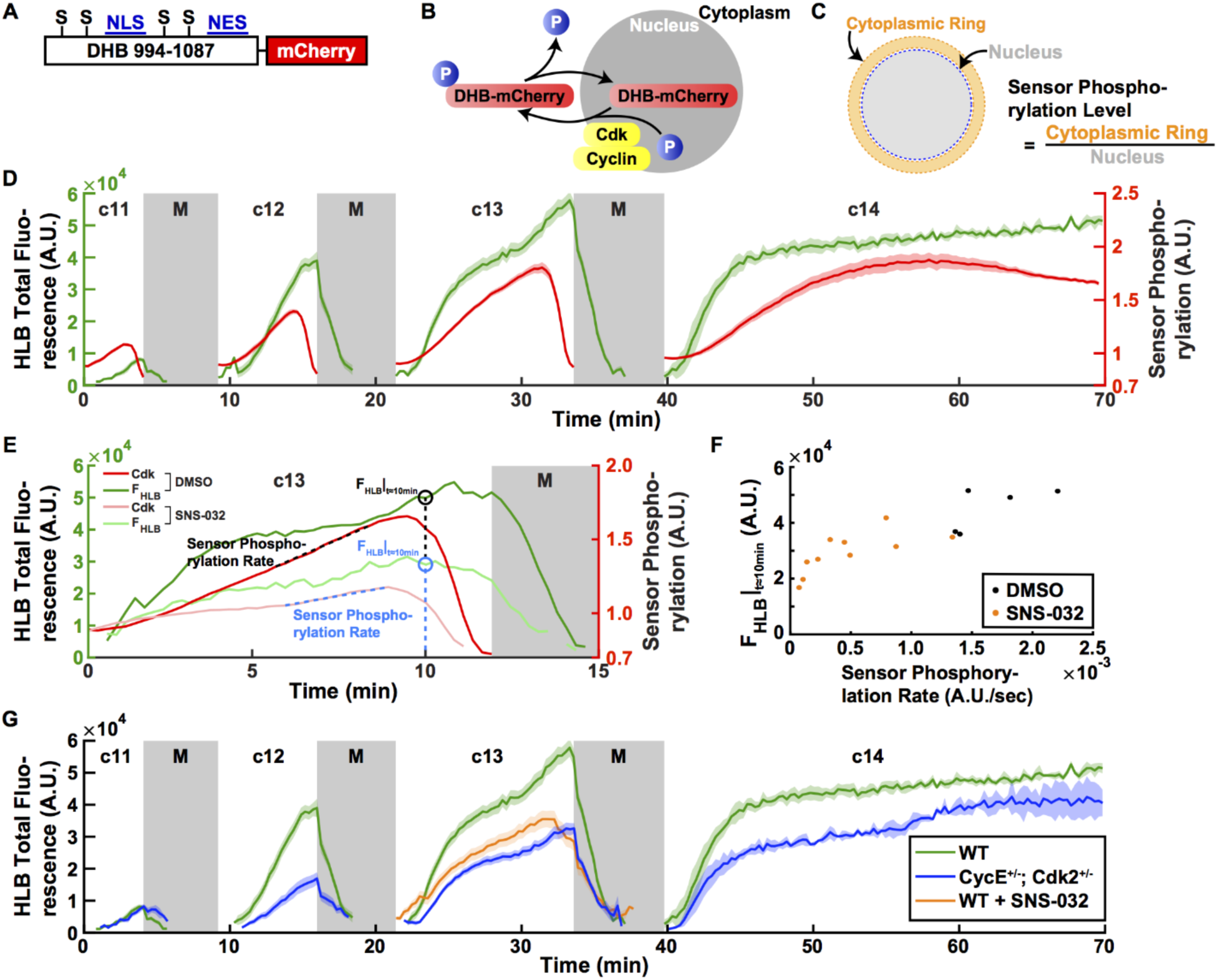
Cdk activity controls the size of HLB. (A) Cdk sensor adapted from Ref (Spencer et al., 2013). Human DNA Helicase B (DHB) amino acids 994-1087 were fused with the fluorescent protein mCherry on the C terminus. S: CDK consensus phosphorylation serine residue, NLS: nuclear localization signal, NES: nuclear export signal. (B) Schematic of Cdk-mediated translocation of the sensor DHB-mCherry. (C) Method used to quantify the sensor phosphorylation level. mCherry fluorescence levels of the cytoplasm surrounding the nucleus (cytoplasmic ring) and the nucleus were measured and the sensor phosphorylation level was estimated by calculating the cytoplasmic-to-nuclear mCherry fluorescence ratio. (D) HLB total fluorescence (green, left axis) and Cdk sensor (red, right axis) dynamics in cycles 11-14. (E) Representative quantifications of HLB and Cdk sensor dynamics for DMSO control and SNS-032 (Cdk2 inhibitor) injected embryos during cycle 13. F_HLB_|_t≈10min_: HLB total fluorescence 10 minutes after injection. Sensor Phosphorylation Rate, quantified as the slope of the sensor readout. (F) HLB total fluorescence plotted against sensor phosphorylation rate. F_HLB_|_t≈10min_: HLB total fluorescence 10 minutes after injection. (G) HLB total fluorescence from embryos laid by WT mothers (green), CycE and Cdk2 double heterozygous mothers (blue), and Cdk2 inhibitor injected embryos (orange).

To further elucidate the role of Cyclin E/Cdk2 and other Cyclin/Cdk complexes in HLB formation, we also altered their activities genetically, using heterozygous mutant mothers (Methods). Embryos laid by mothers carrying heterozygous mutations for both *Cyclin E* and *Cdk2* have smaller HLBs, providing evidence that maternal Cyclin E/Cdk2 activity plays an important role in HLB formation (Figure 5G). In this genetic background, the dynamic of the sensor is almost unperturbed, supporting the idea that the regulation of the biosensor integrates inputs from multiple Cdk complexes (Schwarz et al., 2018) and from cell cycle regulated phosphatases (Figure S5E). Embryos from mothers heterozygous for Cyclin A also showed reduced HLB size, whereas embryos from mothers heterozygous for Cyclin B were similar to wild type embryos (Figure S5G). Altogether, these results argue that Cyclin E/Cdk2 and Cyclin A/Cdk1 modulate HLB formation and may also be indicative of overlapping function of these two kinases (Farrell et al., 2012; Sprenger et al., 1997).

### Cdk activity controls the growth of HLBs by regulating Mxc nuclear concentration

The behavior of HLBs during the nuclear cycles suggested that phosphorylation of Mxc by Cdk complexes may regulate HLB growth (Ma et al., 2000; White et al., 2011; White et al., 2007; Ye et al., 2003; Zhao et al., 1998; Zhao et al., 2000). Therefore, we generated a transgenic fly line expressing a GFP-Mxc in which 22 putative Ser/Pro or Thr/Pro Cdk sites were mutated to Ala, hereafter GFP-Mxc^AP22^ (Figure 6A). We found that GFP-Mxc^AP22^ was not incorporated into HLBs as well as GFP-Mxc, as revealed by the smaller amount of total GFP-Mxc^AP22^ present in foci (Figure 6B). Moreover, GFP-Mxc^AP22^ had a reduced concentration in nuclei compared to GFP-Mxc (Figure 6C, top right panel; Figure 6D). This result suggests that the reduced recruitment to HLB foci might be due to decreased Mxc nuclear concentration and that such concentration might be regulated through phosphorylation. In support of this hypothesis, HLB total fluorescence and nuclear Mxc concentration were strongly correlated (Figures 6B and 6D), and we observed a lower nuclear Mxc concentration in embryos injected with the SNS-032 Cdk2 inhibitor or embryos derived from mothers heterozygous for both *Cyclin E* and *Cdk2* (Figure 6B). In all genetic conditions altering *Cyclin E, Cdk2* and/or *Cyclin A*, the Mxc concentration within the HLB remains unaltered (Figure 6E), arguing that changes in cell cycle control do not change the phase diagram of HLB formation. Moreover, the relationship linking HLB size to Mxc nuclear concentration is quantitatively the same for all genetic conditions (Figure 6F). Collectively, these results argue that Cdk-dependent phosphorylation regulates the nuclear level of Mxc, which in turn controls HLB growth. This interpretation is strengthened by analysis of HLB dynamics in embryos injected with Roscovitine, a small molecule inhibitor of Cdk1 and Cdk2. Inhibition of Cdk activity results in dephosphorylation of the biosensor and arrest of the nuclear cycle, as expected (Figure S6C). However, HLBs persist since Mxc nuclear levels are high enough for phase separation (Figures S6A and S6B). Thus, we conclude that Mxc can form HLBs even in the absence of Cdk-dependent phosphorylation, as long as its concentration is sufficiently high, an observation also consistent with previous findings showing that HLBs form in G1 when Cdk1 and Cdk2 activities are absent and HLBs contain unphosphorylated Mxc (Liu et al., 2006; White et al., 2011).

**Figure 6.**
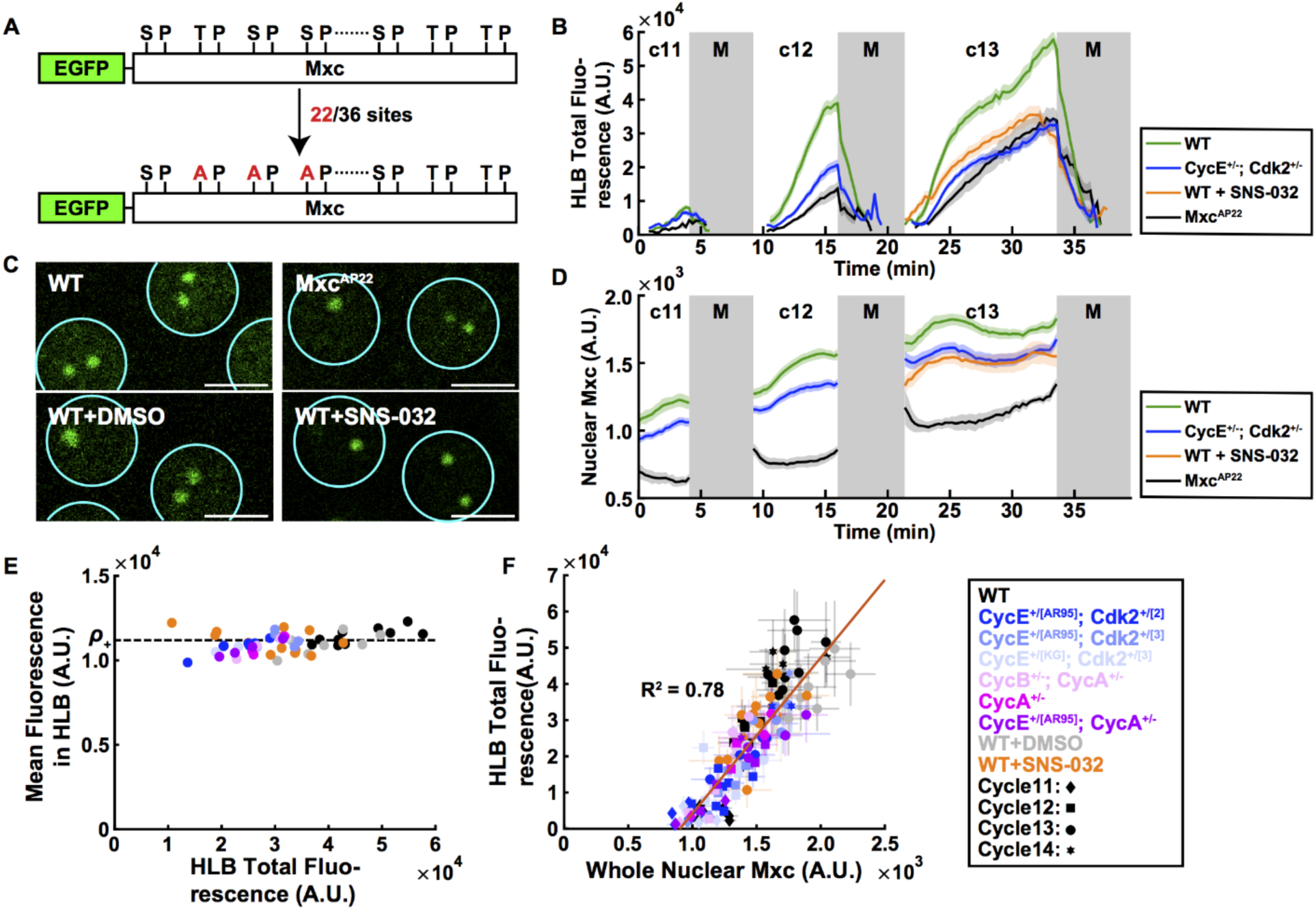
Nuclear Mxc concentration is regulated by Cdk activity and determines HLB size. (A) Schematic of the non-phosphorylateable mutant Mxc (Mxc^AP22^). Out of the 36 consensus SP or TP Cdk phosphorylation sites found in the Mxc protein, numbers 2-23 throughout the N-terminus were mutated to alanines. (B) HLB total fluorescence for embryos laid by WT (green), Cyclin E/Cdk2 double heterozygotes (blue), Mxc^AP22^ (black) mothers, and Cdk2 inhibitor injected (orange) embryos. (C) Confocal images of HLBs for WT (top left), GFP-Mxc^AP22^ expressing embryos (top right), DMSO control injected embryos (bottom left), and Cdk2 inhibitor injected embryos (bottom right). (D) Nuclear Mxc concentration for WT (green), Cyclin E/Cdk2 double heterozygotes (blue), Cdk2 inhibitor injected (orange), and Mxc^AP22^ embryos (black). (E) Mxc concentration inside HLBs for various genetic conditions altering cell cycle regulators. (F) The simple dependency of HLB size vs Mxc concentration (From Figure 3K) still holds with different genetic conditions. A.U., arbitrary units. c, cycle. M, mitosis. Scale bars, 5µm.

### Reducing Cdk2 activity and HLB size results in misprocessing of nascent histone mRNAs

To test whether the mechanisms controlling HLB growth are functionally linked to histone mRNA biogenesis, we determined whether perturbations of Cdk2 activity affect histone pre-mRNA processing. RD histone mRNAs are not polyadenylated, and instead end in a 3’ stem loop that binds the stem loop binding protein (SLBP) (Marzluff et al., 2008). SLBP is required for the endonucleolytic event resulting in this unique mRNA 3’ end (Sullivan et al., 2001) (Figure 7A, top panel). In *Drosophila Slbp* mutants, histone pre-mRNA processing is impaired and RNA polymerase continues transcribing downstream of the normal processing site, resulting in longer nascent transcripts (Sullivan et al., 2001). Consequently, the efficiency of histone pre-mRNA processing can be measured by *in situ* hybridization using a probe complementary to the region downstream of the histone H3 pre-mRNA processing site (Lanzotti et al., 2002) (Figure 7A, bottom panel). Detection of focal, nuclear signal with this probe indicates the presence of nascent mRNA that has not been cleaved at the normal location. In embryos derived from *Slbp*^*+/-*^ heterozygous mothers, the level of nascent, unprocessed H3 mRNA is increased compared to wild type, indicating less efficient pre-mRNA processing (Figure 7B; Figure S7B). Using this approach, we found that the level of nascent, unprocessed H3 mRNAs is elevated in embryos laid by mothers heterozygous for both *Cyclin E* and *Cdk2* (Figure 7C; Figure S7B), suggesting that reduced HLB size impairs histone pre-mRNA processing. We propose that the amount of processing enzymes recruited to HLBs is proportional to their size and that the tight control of HLB growth ensures the efficient processing of nascent histone mRNAs at the maternal-to-zygotic transition.

**Figure 7.**
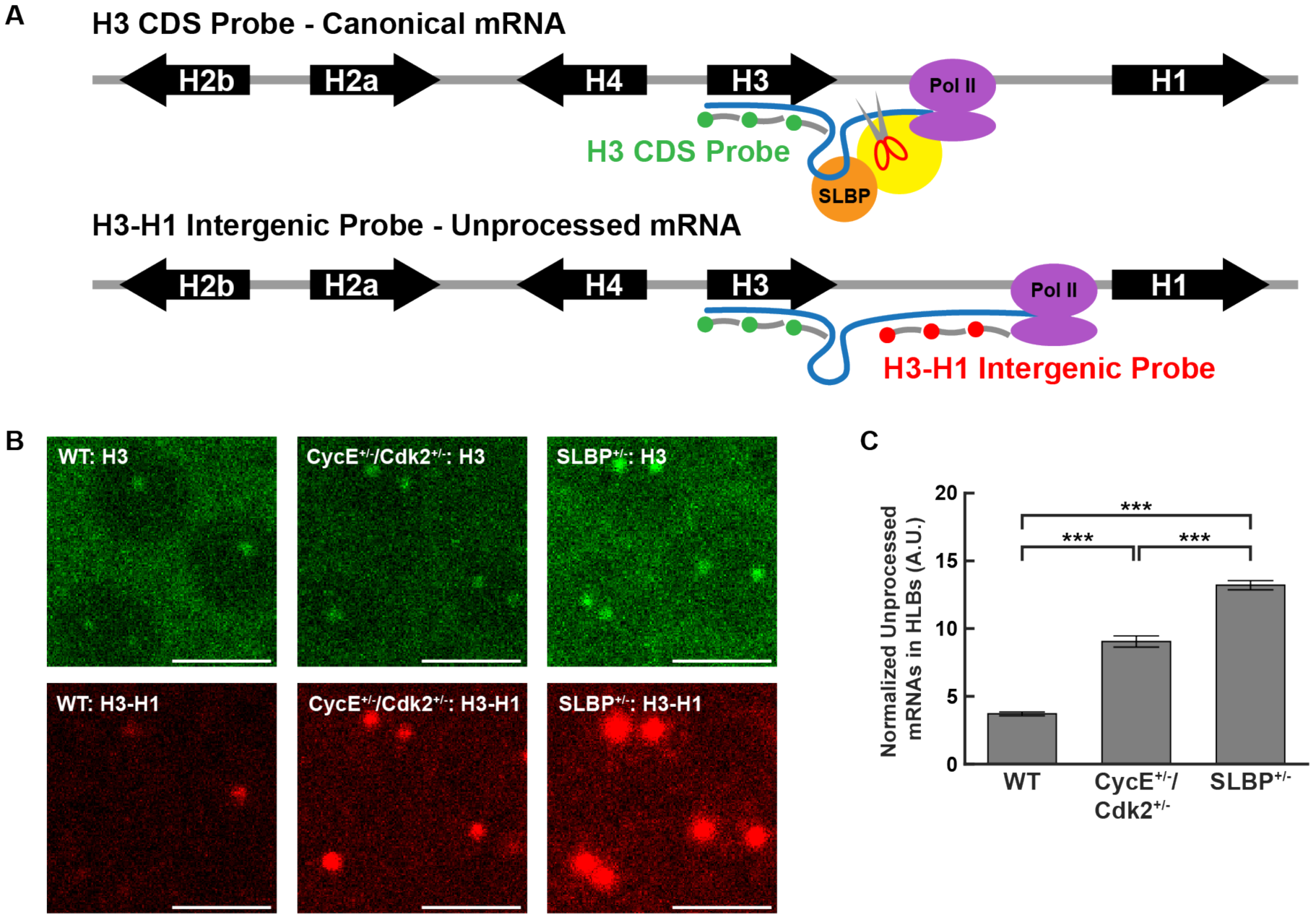
HLB size determines its processing ability. (A) Schematic of the fluorescence in situ hybridization (FISH) probe for histone H3 coding sequence (CDS) (top) and H3-H1 intergenic sequence (bottom), to detect the level of nascent transcription of canonical mRNA and misprocessed mRNA, respectively. Yellow blob is the cleavage complex. (B) Representative images of fixed embryos stained with the H3 CDS probe and H3-H1 intergenic probe, for embryos laid by WT, CycE^+/-^/Cdk2^+/-^, and SLBP^+/-^ mothers. Scale bars, 5µm. (C) Quantification of normalized misprocessed mRNA levels for embryos laid by WT, CycE^+/-^/Cdk2^+/-^, and SLBP^+/-^ mothers. Error bars, SEM. ***, p<0.001.

## DISCUSSION

In this study, we investigated the mechanisms of formation of the HLB, an evolutionarily conserved nuclear body that controls the production of RD histone mRNAs. We demonstrated that HLB formation is driven by two mechanisms: 1) seeding from the histone locus, which is enhanced by transcription and initiates liquid-liquid phase separation; and 2) nuclear accumulation of Mxc, which is controlled by Cdks. Zygotic expression of RD histone genes and the concentration of Mxc together determine the rate of growth of HLBs. Collectively, our findings reveal a strategy for the regulation of the growth of nuclear bodies: seeding by genomic loci and transcription overcomes the stochastic step of nucleation and control of nuclear levels by signaling ensures accurate growth dynamics. The importance of seeding by RNA to overcome the stochasticity of nucleation was previously demonstrated for the formation of the nucleolus (Falahati et al., 2016), arguing that seeding represents a general strategy for the control of the formation of nuclear bodies. Our theoretical and experimental work further extends this concept by demonstrating that in an out-of-equilibrium regime the level of seeding contributes to determine the growth rate of nuclear bodies, demonstrating an additional layer of regulation. The concentration of nucleolar components determines the size of the nucleolus (Weber and Brangwynne, 2015), similarly to our observations for the control of the HLB by nuclear Mxc concentration. In the future, it will be interesting to determine which signaling mechanisms control nucleolar component concentration in the nucleus, and whether there is a relationship between size and function similar to the one we have uncovered here for the HLB.

### The HLB is a phase separated nuclear body

Phase separation is a highly stochastic process when it is initiated by the random formation of droplets by nucleation. In such a scenario, fluctuations must drive the formation of structures bigger than the critical radius needed to overcome the effects of surface tension. As a consequence, the size of phase-separated objects during their growth phase can be quite variable. Mechanisms of seeding can provide a way to accurately control the initial step of phase separation, thus reducing variability and ensuring control of the number of phase-separated objects (Falahati and Wieschaus, 2017). Indeed, the HLB invariably forms at RD histone loci, as clearly illustrated by the syncytial fly embryo in which all nuclei only have either one or two HLBs depending on whether the homologous RD histone loci are paired or not. In the absence of histone loci, Mxc molecules can still phase separate, but the resulting foci are variable in size and number. Our theoretical analysis also shows that in the initial, non-equilibrium phase of growth the levels of seeding not only overcome the need for nucleation but, together with nuclear concentration, directly impact the rate of growth of phase-separated objects. Thus, our experimental and theoretical analysis argue for a non-equilibrium mechanism that allows the precise control of the size of a nuclear body.

Our data presented here and previously (Salzler et al., 2013) indicate that transcription of RD histone genes plays a crucial role in facilitating the assembly of the HLB. In addition, tethering an engineered H2b mRNA to a *lacO* array integrated into the HeLa cell genome drives the recruitment of HLB factors, including NPAT, and formation of ectopic HLBs (Shevtsov and Dundr, 2011). These observations suggest that interactions between nascent mRNA and proteins forming the HLB might facilitate phase separation. Consistent with this idea, recent studies have indicated that changes in RNA/RNA-binding protein ratios or mRNA structure can modulate the formation and composition of liquid-liquid phase-separated bodies (Langdon et al., 2018; Maharana et al., 2018). The specific nature of nascent RD histone RNA/protein interactions and their role in the establishment of the HLB remains to be fully elucidated. Importantly, during later developmental stages, HLBs are present in cells that are not replicating (Liu et al., 2006; White et al., 2007), arguing that the histone locus is also able to seed the formation of HLBs in the absence of transcription (RD genes are transcribed only in the S phase of the cell cycle). These results are consistent with our alpha-amanitin injections in which HLB size is highly reduced but still only one or two HLBs are observed (likely at the histone locus). These observations suggest that components of the HLBs must be able to bind to both DNA and nascent mRNA and that high transcriptional levels increase these binding interactions, thus enhancing the ability of the histone locus to seed the formation of HLBs. In the future, it will be important to elucidate which protein-DNA and protein-mRNA interactions are important for the regulation of HLB growth.

Carrying out RD histone mRNA transcription and processing in a phase-separated compartment within the nucleus may facilitate proper histone mRNA biosynthesis. Several of the factors involved in canonical mRNA polyadenylation, including those that carry out pre-mRNA cleavage (e.g. Symplekin and the CPSF73/CPSF100 endonuclease), are also required for formation of the unique metazoan RD histone mRNA 3’ end (Marzluff and Koreski, 2017). The cleavage complexes involved in polyadenylation and RD histone mRNA processing are distinct (Sullivan et al., 2009), and thus phase transition might function to exclude the polyadenylation machinery from accessing RD histone genes. In addition, phase transition might also function to drive a high local concentration of cleavage factors, thereby increasing the efficiency of RD histone mRNA biosynthesis (Tatomer et al., 2016; Wagner et al., 2007).

### HLB size contributes to efficient RD histone mRNA biosynthesis

In addition to the biophysical properties afforded by phase transition, the size of nuclear bodies may contribute to their function. Our microscopic and immuno-FISH data reveal that HLB size in the early fly embryo is remarkably consistent and directly related to levels of nascent RD histone mRNAs. These observations suggest tight control of HLB size, and thus perhaps that HLB size is functionally relevant. Indeed, we found that reducing HLB size results in the appearance of misprocessed nascent H3 mRNA. Histone genes are among the most highly transcribed, and thus a large HLB might be needed to ensure sufficient amounts of factors are concentrated to process the many RNAs being synthesized from the up to 1000 potentially active genes within the RD histone gene array. HLB size is linked to the nuclear concentration of Mxc, which we found depends on the activity of Cdks. Our data with a mutant version of GFP-Mxc lacking 22 Cdk consensus sites suggests that this regulation likely occurs at least in part by direct phosphorylation of Mxc by Cyclin E/Cdk2. We therefore propose that control of HLB size via CDK activity is an important mechanism by which *Drosophila* embryos achieve functional control of histone mRNA biogenesis.

In conclusion, our work identifies the molecular and physical mechanisms of formation of the Histone Locus Body, a nuclear membrane-less organelle playing a crucial role in the biosynthesis of RD histone mRNAs. Our results indicate that reducing HLB size impairs histone pre-mRNA processing, thus providing a direct link between size and function of a nuclear body. Our HLB work suggests a general strategy for accurate size control of nuclear bodies forming by phase-separation centered on a mechanism integrating transcription and kinase signaling. We speculate that the coupling of seeding from nascent mRNAs and signaling-dependent regulation of nuclear concentration represents a general mechanism ensuring that nuclear bodies provide accurate control of gene regulatory functions.

## Acknowledgments

We thank the Bloomington Drosophila Stock Center, the Kyoto Drosophila Stock Center for providing stocks. We thank the Drosophila Genomics Resource Center for constructs. We thank Anna Chao and James Kemp for help with experiments. We acknowledge discussions with James Kemp and Daniel Lew. We thank Michel Bagnat, Sharyn Endow, Brigid Hogan, Bill Marzluff, Bernard Mathey-Prevot, Chris Nicchitta and members of the Di Talia lab for comments on the manuscript. This work was supported by the E. Bayard Halsted Fellowship (to W.H.), Schlumberger Faculty for the Future Fellowship and an HHMI International Student Research Fellowship (to V.E.D.), and NIH (R01-GM058921 to R.J.D. and R01-GM122936 to S.D.T.).

## Author Contributions

Conceptualization: W.H., R.J.D. and S.D.T; Methodology: W.H., M.T., V.E.D., E.A.T., R.J.D. and S.D.T.; Software: W.H., M.T. and S.D.T.; Validation: W.H. and M.T.; Formal Analysis: W.H., M.T. and S.D.T.; Investigation: W.H. and M.T.; Resources: W.H., M.T., V.E.D., E.A.T., R.J.D. and S.D.T.; Data Curation: W.H.; Writing-Original Draft: W.H., R.J.D. and S.D.T.; Writing-Review & Editing: W.H., M.T., V.E.D., E.A.T., R.J.D. and S.D.T.; Visualization: W.H.; Supervision: R.J.D. and S.D.T.; Project Administration: S.D.T; Funding Acquisition: S.D.T.

## METHODS

### Molecular Biology and Transgenic Flies

The DHB-mCherry Cdk sensor (human DNA Helicase B amino acids 994–1087 fused with mCherry on the C-terminus) was cloned in the plasmid pBabr containing the maternal Tubulin promoter and the spaghetti squash 3’ UTR (a gift of Yu-Chiun Wang and Eric Wieschaus, Princeton University), using Gibson assembly (NEB). The mutant version of Cdk sensor DHB-mCherry^4A^ (Figure S5) was generated by introducing mutation of 4 putative phosphorylation serine residues into alanines (S12A, S28A, S55A, S65A). The pBabr construct was then injected into ϕC31 based fly line zh-86Fb (http://www.flyc31.org), which has the insertion site at 86Fb on the 3^rd^ chromosome.

GFP-Mxc^AP22^ fragment was generated by mutagenizing serines and threonines of 2^nd^-23^rd^ putative phosphorylation sites (Ser/Pro or Thr/Pro) from the N-terminus to alanines (Ala) on the *mxc* gene (GenScript). The first phosphorylation site was left out as it is in the self-interacting LisH domain. The synthesized DNA fragment, encompassed between the unique intrinsic restriction enzyme sites MreI and ScaI and containing the 22 mutagenized phosphorylation sites, was cloned into a pENTR vector (ThermoFisher Scientific) with full-length *mxc* cloned in it, by replacing the N-terminal fragment containing the wild-type phosphorylation sites. *gfp-mxc*^*AP22*^ transgenic fly line was generated by injecting *y*^*1*^*w*^*1118*^;; PBac[*y*^*+*^-attP-3B]VK00033 embryos (BestGene, Inc.) with a ϕC31-compatible vector containing an N-terminal GFP tag and expression of the fusion protein was driven by the *ubiquitin* promoter in pUGW.

All live imaging experiments were done with embryos expressing transgenic Mxc-GFP in the presence of endogenous Mxc. For convenience, these Mxc-GFP lines were indicated as ‘WT’ in the main text and figures. All immuno-FISH experiments were done with embryos that do not express any transgenic Mxc. The ‘WT’ embryos in these experiments were *w*^*1118*^. Details on all transgenic flies used in this study are listed in the Key Resources Table.

### Live imaging of HLBs

Flies of desired genotype were placed in a cage with an apple juice agar plate and yeast paste. Embryos that were 0-2 hours old were collected and dechorionated by bleaching for 1-1.5 minutes in 50% bleach. Dechorionated embryos were then mounted on an air-permeable membrane with halocarbon oil 27 (Sigma-Aldrich, CAS Number 9002-83-9) and covered with a cover slip. Images were acquired through confocal microscope Leica SP8 and its software Leica Application Suite X (LAS X), with 600Hz scan speed, 20× magnification objective with immersion oil (HC PL APO CS2 20×/0.75 IMM), and 10× zoom which gave 0.073µm/pixel resolution. 488nm laser was used to excite GFP-Mxc, and 561nm laser was used to excite the DHB-mCherry Cdk2 sensor. HyD detectors were used for both channels.

### Immuno-FISH

#### FISH Probes

We designed FISH probes targeted for *Drosophila* histone H3 coding sequence to measure the amount of nascent histone transcripts (Figure 7A, top panel). We also designed similar probes for the H3 downstream sequence to stain for the misprocessed mRNAs^15^ (Figure 7A, bottom panel). Custom Stellaris® FISH Probes were designed utilizing the Stellaris® RNA FISH Probe Designer (Biosearch Technologies, Inc., Petaluma, CA) available online at www.biosearchtech.com/stellarisdesigner (Version 4.2). The histone H3 coding sequence probes were hybridized with the green fluorescent dye Fluorescein, and the histone H3-H1 intergenic sequence probes were hybridized with the far-red fluorescent dye Quasar® 670 Dye.

#### Fixation

Dechorionated embryos were suspended in embryo fixation solution (8mL nuclease-free water, 1mL 10x PBS, 1mL 37% formaldehyde concentrate, and 10mL heptane) and were shaken for 20 minutes in a scintillation vial. Bottom aqueous phase was removed, 10mL methanol was added, and the vial was shaken vigorously for 30 seconds. Devitellinized embryos were pipetted out from the bottom of the methanol layer. The embryos were washed in fresh methanol for 3 times, transferred to a new tube, and were stored at - 20°C before staining.

#### Staining

Fixed embryos were quenched by rocking in PBT (0.1% Tween-20 in 1x PBS) for 10 minutes for 3 times, rocked in 50:50 PBT:wash buffer (2x SSC, 10% formamide) for 10 minutes, and rocked in the wash buffer for 2 times for 5 minutes at room temperature. Wash buffer was removed and 500mL hybridization buffer (1g dextran sulfate in 10mL wash buffer) was added and incubated for 2 hours at 37°C for pre-hybridization to prevent nonspecific binding. FISH probe mixture was made by diluting the probes to 50nM in the hybridization buffer. The pre-hybridization mixture was removed, the probe mixture was added, and was incubated in the dark at 37°C overnight. The probe mixture was then replaced with blank hybridization buffer and incubated at 37°C in the dark for 30 minutes. The embryos were then washed in the wash buffer for 4 times for 15 minutes, 3 times in PBT for 10 minutes in the dark. Primary antibody mixture (1:20000 MPM-2 in PBT) was added and rocked at 4°C in the dark overnight. The embryos were washed in PBT 3 times for 10 minutes in the dark. Secondary antibody mixture (1:500 Goat anti-Mouse IgG (H+L) Cross-Adsorbed Secondary Antibody, Alexa Fluor 568 in PBT) was added and rocked at room temperature for 2 hours. The embryos were washed in PBT 3 times for 10 minutes and were mounted onto the slides with the Aqua-Poly/Mount Coverslipping Medium.

### Injection

Following the dechorionation, the embryos were aligned and glued to a cover slip with a double-sided tape. The embryos were desiccated in an airtight glass container filled with the desiccant Drierite (https://secure.drierite.com/) for 6-8 minutes. About 4μl Inhibitor solution was loaded to the needle Femtotip® II. Using Eppendorf® FemtoJet® Microinjector, embryos were then injected. Embryos were immediately covered with halocarbon oil 700 (Sigma-Aldrich, CAS Number 9002-83-9) following the injection and imaged.

## QUANTIFICATION AND STATISTICAL ANALYSIS

### Quantitative image analysis

Confocal images were exported as .tif files from LAS AF Lite software and were imported into MATLAB for image analysis. Different custom scripts were written for different analyses performed. HLB masks were generated using threshold-based segmentation and HLB size was approximated by measuring the space occupied by multiple segmented masks in different z-planes (Figure 1B). The total fluorescence of HLBs were then calculated by multiplying the approximated size to the average fluorescence level in the HLB masks. The HLB total fluorescence throughout this study was quantified by summing the total fluorescence per nucleus, as two HLBs on the two different homologous chromosomes sometimes fuse with each other and this fusion resulted in simple addition of the two droplets as opposed to promoting further growth (Figure 1F). The Cdk sensor phosphorylation level was quantified by measuring the average DHB-mCherry fluorescence levels in the nucleus and the surrounding cytoplasm respectively, and then calculating the cytoplasmic-to-nuclear ratio. The circularity index of HLB during fusion was calculated by 4π*A/C*^2^ where A is the area of the HLB mask and C is the perimeter of the HLB mask. All statistical analyses were performed by running a one-way ANOVA followed by a Turkey’s Pair test to compare two sets of data.

### Analysis of HLB FRAP experiments

Since HLBs can grow significantly within the time scale of fluorescence recovery after photobleaching, we used a mathematical model to estimate the export rate *k*_*off*_. Our model assumed that the import rate *k*_*on*_ is independent of the concentration inside the HLB, an assumption supported by the observation that the concentration of nucleoplasmic Mxc does not vary significantly over an S phase. By denoting *C*_u_(t) and *C*_b_(t) the concentrations of unbleached and bleached HLBs respectively, we obtain:

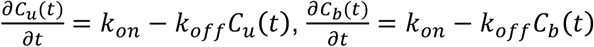

Subtracting the second from the first equation, we get

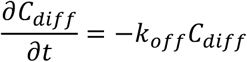

where *C*_*diff*_ (t) *= C*_u_(t) *-C*_b_(t) While *k*_*off*_ depends on time (see correlation with HLB size), we assume that on the timescale of the FRAP experiments *k*_*off*_ is essentially constant. Thus, it can be expressed as:

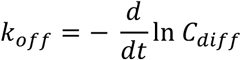

This implies that we can estimate *k*_*off*_ from the inverse slope of a linear fit of ln *C*_*diff*_ as a function of time.

### Quantification of Immuno-FISH results

The number of total histone transcripts and the number of total phospho-Mxc were estimated by measuring the total fluorescence from each staining. Total fluorescence was acquired with similar methods used for live imaging of HLB from the previous section. Normalized misprocessed mRNAs level in HLBs was calculated by total fluorescence of H3-H1 intergenic probe divided by that of H3 CDS probe. Statistical analysis was performed by running a one-way ANOVA followed by a Turkey’s Pair test. n=409, 343, and 681 HLBs respectively for WT, CycE^+/-^/Cdk2^+/-^, and SLBP^+/-^.

### Statistical analysis

All statistical analyses were performed using JMP Pro. Statistical comparisons between multiple samples was by one-way ANOVA followed by Tukey’s test to compare all pairs. For all measurements, at least three biological replicates were used unless otherwise noted.

