## Supplementary Material for "CDK-regulated phase separation seeded by histone genes ensures precise growth and function of Histone Locus Bodies"

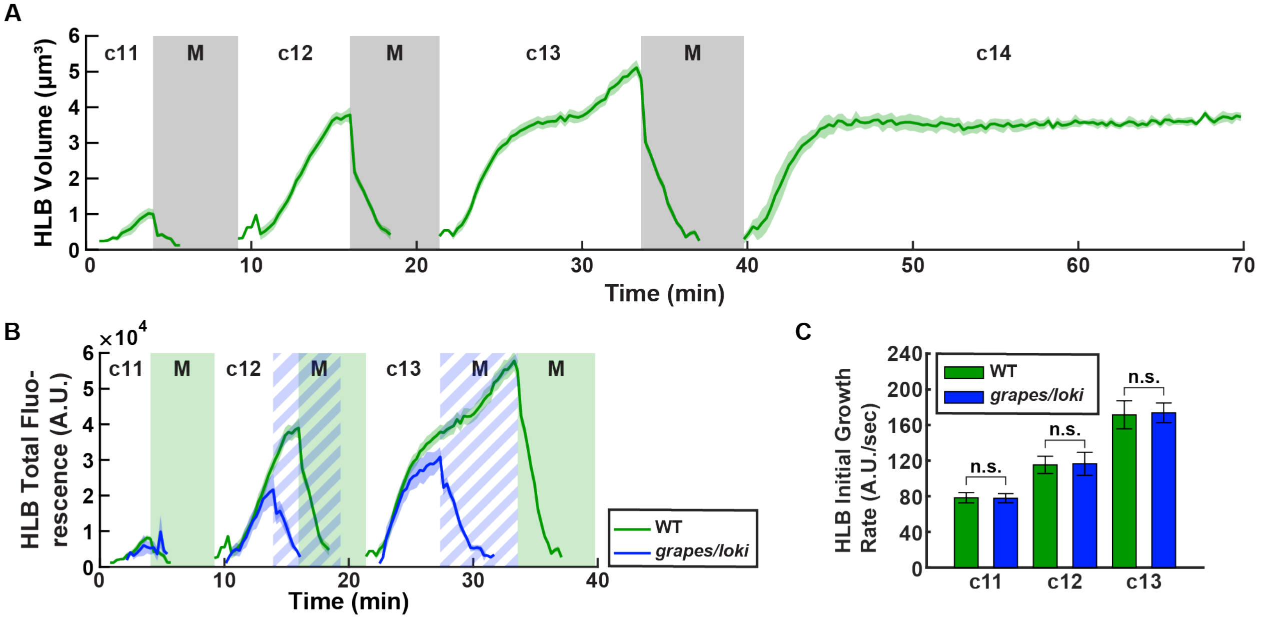


**Figure S1. Detailed analysis of HLB dynamics**

(A) HLB volume during cycles 11-14 for WT embryos. (B) HLB total fluorescence during cycles 11-13 for the *grapes/loki* (mammalian ortholog of *Chk1/Chk2*) mutants. The beginning of S phase is aligned with that of WT for comparison, as this mutant has a shorter S phase. Mitoses for the *grapes/loki* mutants are marked as striped blue area. (C) The initial growth rate of HLB for WT and *grapes/loki* mutants measured by slopes of Figure S1D. Error bars, standard deviation. n.s., not significant. A.U., arbitrary units. c, cycle. M, mitosis.

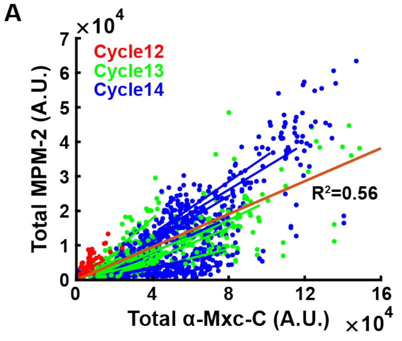


**Figure S2. MPM-2 staining correlates with Mxc during S phase**

(A) Total fluorescence from the MPM-2 antibody staining and the Mxc antibody staining. Orange line is the best fit line through the origin. Each colored line is the best fit line for each embryo, in the color of its respective cycle. A.U., arbitrary units.

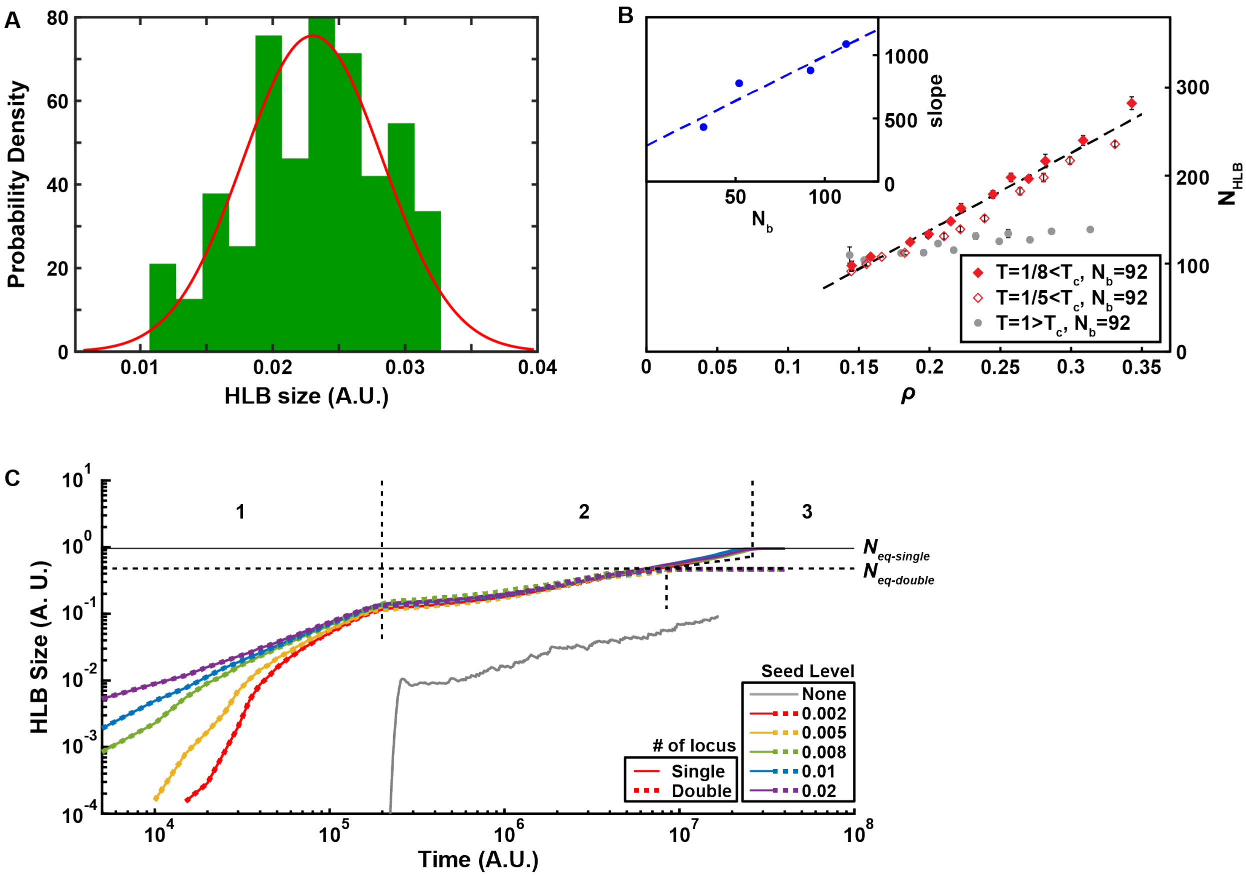


**Figure S3. Additional simulation results**

(A) HLB size distribution from a simulation that includes seeding. (B) Average volume of the HLB droplet, $N_{HLB}$, as a function of the average concentration outside the droplet $\rho$, after $10^5$ MC steps of the model (6) on a cubic lattice of $L = 43$ sites with seeding level $N_b = 92$. Filled diamonds correspond to $T = 1/8$. Dashed lines are the linear fts of the data as $N_{HLB} = \alpha\rho+\beta$. Empty diamonds correspond to a slightly higher temperature, but still below $T_c$, $T = 1/5$. Gray filled circles correspond to $T = 1$, slightly above $T_c$. Inset: Slope $\alpha$ obtained from the linear fit of the main panel as a function of $N_b$. The main figure was $N_b = 92$, and additional points of $N_b = 32$, $N_b = 52$, and $N_b = 112$ are shown. (C) HLB growth simulation through numerical solution of Cahn-Hilliard (CH) equation in 2D lattice, with single seed locus (solid line) and double seed loci (dashed line), with different levels of seed (same as Figure 3I). $N_{eq\text{-}single}$, equilibrium size for the single seed locus simulation. $N_{eq\text{-}double}$, equilibrium size for the double seed loci simulation. A.U., arbitrary units. c, cycle. M, mitosis.

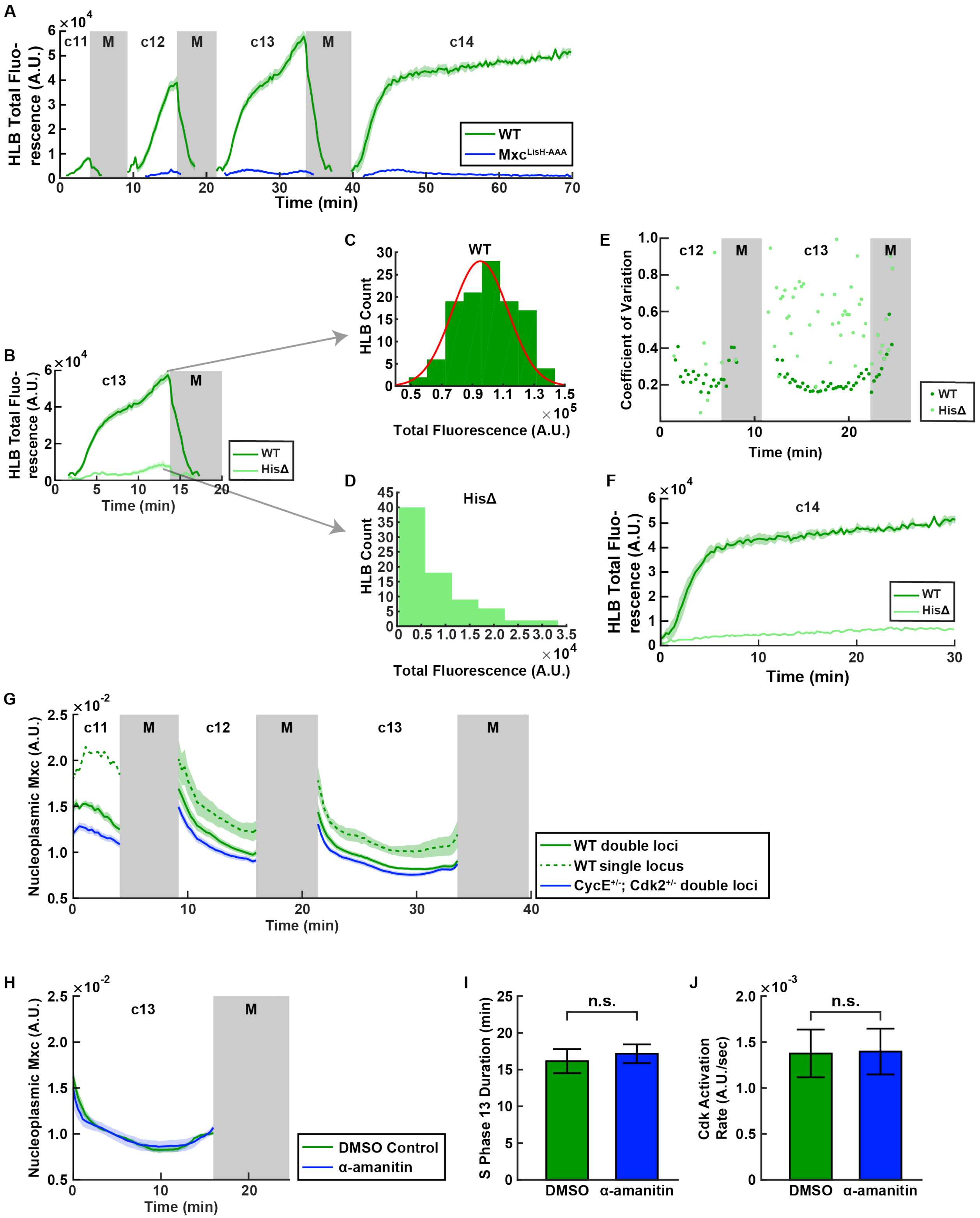


**Figure S4. Additional data on $Mxc^{LisH\text{-}AAA}$ and transcriptional seeding**

(A) HLB total fluorescence during cycle 11-14 for WT and *mxc*$^{LisH\text{-}AAA}$ mutants. (B) HLB total fluorescence for WT and histone deficiency (HisΔ) embryos for cycle 13 (from Figure 4B). Arrows indicate representative examples of HLB total fluorescence distribution for (C) WT and (D) HisΔ.

(E) Comparison of HLB total fluorescence coefficient of variation (CoV) for WT and HisΔ embryos. (F) Mxc-GFP total fluorescence for WT and HisΔ embryos for cycle 14. Longer S phase allows for proto-HLBs to grow to a more significant size. (G) GFP-Mxc level in the nucleoplasm over cycles 11-13 for different genotypes. (H) GFP-Mxc level in the nucleoplasm over cycle 13 for DMSO control and α-amanitin injected embryos. (I) Cycle 13 S-phase duration for DMSO control and α-amanitin injected embryos. (J) Cdk sensor phosphorylation level for DMSO control and α-amanitin injected embryos. A.U., arbitrary units. c, cycle. M, mitosis.

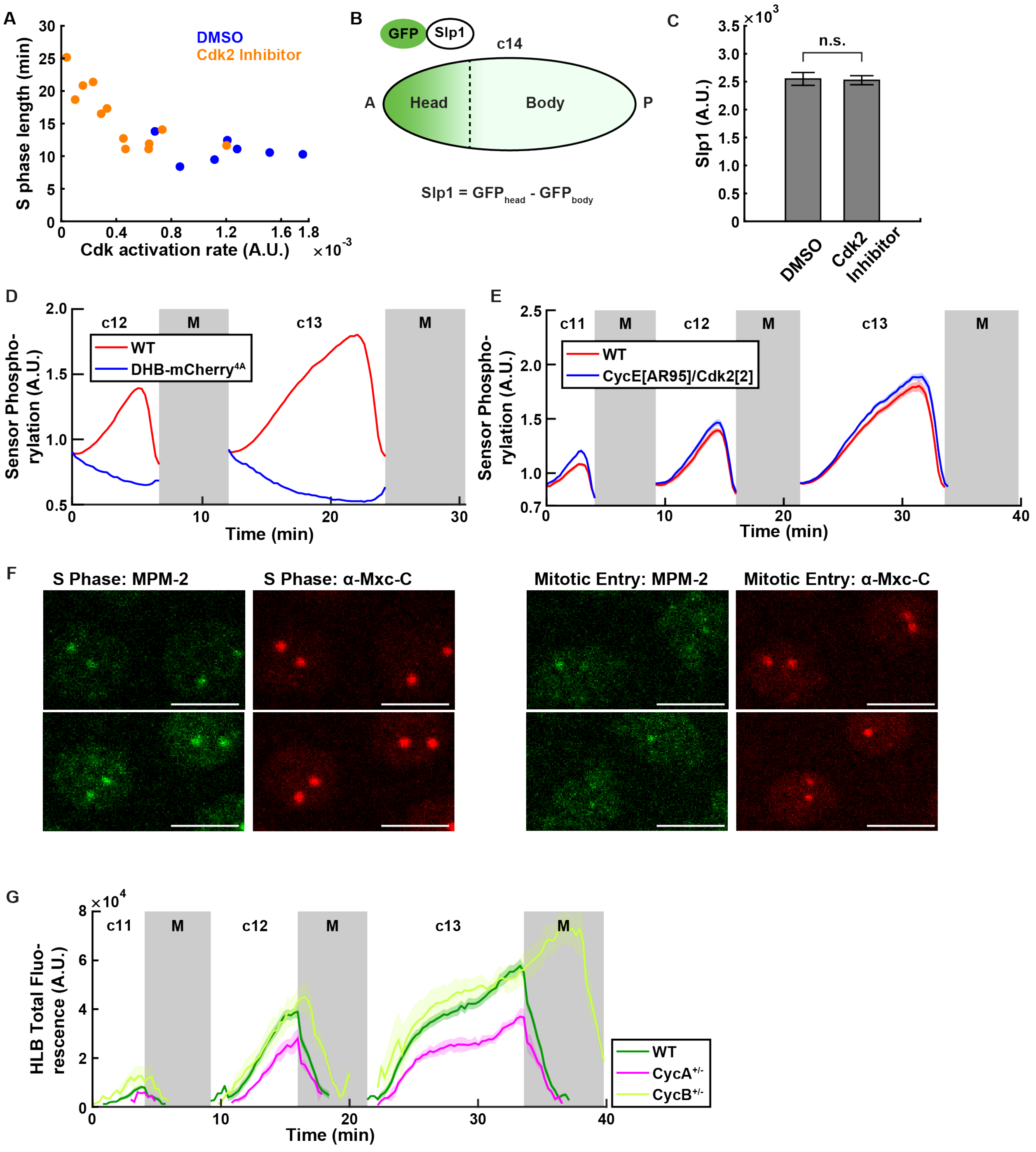


**Figure S5. Regulation of S-phase Cdk phosphorylation and its effect on HLB growth**

(A) S phase length and Cdk activation readout from the Cdk sensor. (B) To investigate whether injections of SNS-032 inhibitor alter transcription by acting on Cdks other than Cdk2, we quantified the expression levels of the transcription factor sloppy paired 1 (Slp1), a zygotically expressed gene, during cycle 14. We compared the GFP-Slp1 protein for DMSO control and SNS-032-injected embryos over a period of about 4.2 minutes during cycle 14. (C) Slp1 level. Error bars, standard deviation. n.s., not significant. (D) Cdk activity measured from WT sensor (red) and the mutant sensor DHB-mCherry$^{4A}$ (blue). (E) Cdk activity measured from the sensor for embryos laid by WT mothers (red) and CycE$^{+/-}$; Cdk2$^{+/-}$ mothers (blue). (F) MPM-2 and an antibody against Mxc

C-terminus (α-Mxc-C) staining for nuclei in S phase (left) and mitotic entry (right). The S phase staining for MPM-2 and α-Mxc-C correlate strongly (see also Figure S2 for a quantification). However, as the nuclei enter mitosis, MPM-2 foci have disappeared or have become smaller while Mxc remains inside the HLB. The decrease in MPM-2 fluorescence preceding mitosis mirrors the decrease in activity readout from the biosensor in (D), suggesting that Mxc and the DHB peptide used in the biosensor are phosphorylated/dephosphorylated with similar dynamics during the cell cycle. Scale bars, 5μm. (G) HLB total fluorescence for embryos laid by WT (green), $CycA^{+/-}$ (magenta), and $CycB^{+/-}$ (lime) mothers. For comparison, the beginning of S phase for $CycB^{+/-}$ embryos was aligned with other genotypes even if they had a longer S phase.

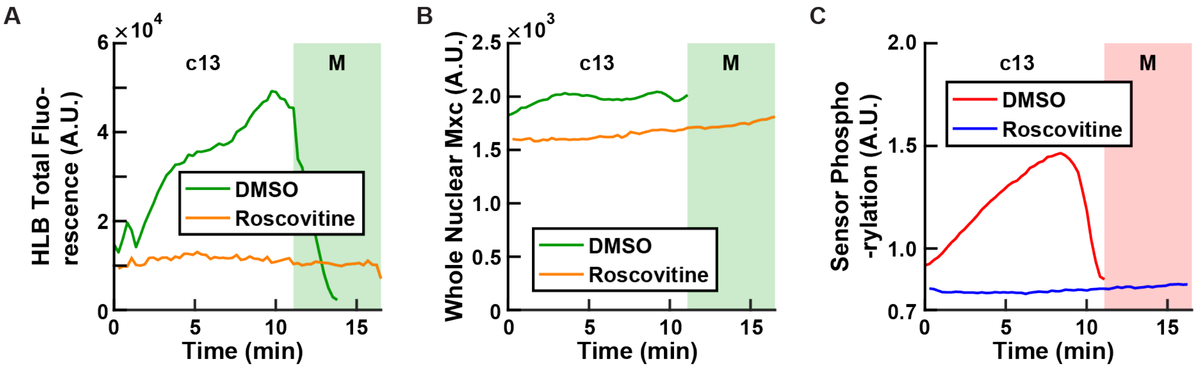


**Figure S6. Phase separation of Mxc is dependent on its nuclear concentration**
(A) HLB total fluorescence during cycle 13 for DMSO control and roscovitine injected embryos. (B) Nuclear Mxc concentration during cycle 13 for DMSO control and roscovitine injected embryos. (C) Cdk sensor phosphorylation level during cycle 13 for DMSO control and roscovitine injected embryos. Notice how roscovitine injected embryos have very low readout from the biosensor (C), are arrested in interphase with a lower concentration of Mxc (B), which is still above threshold so that small HLB persist in the absence of Cdk activity (A). A.U., arbitrary units. c, cycle. M, mitosis.

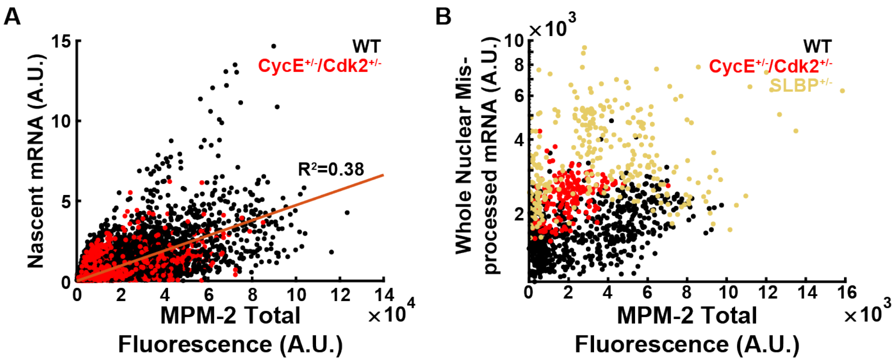


**Figure S7. Additional FISH results**
(A) Levels of nascent mRNA measured from the H3 CDS probe and MPM-2 total fluorescence, for embryos laid by WT (black) and $CycE^{+/-}$; $Cdk2^{+/-}$ (red) mothers. Orange line is the best fit line through the origin. (B) Levels of total misprocessed mRNA level in the nucleus and MPM-2 total fluorescence for WT (black), $CycE^{+/-}$; $Cdk2^{+/-}$ (red) mothers and $SLBP^{+/-}$ (yellow). A.U., arbitrary units.
